# Reconstructing life history of the 18^th^ century priest from Prozorje, Croatia: bioarchaeological and biochemical approaches

**DOI:** 10.1101/2025.02.04.636416

**Authors:** T. Kokotović, M. Carić, S. Stingl, M. Novak, J. Belaj

## Abstract

The study presents a case of the adult male individual dated to the Early Modern Period buried at the church of St. Martin at Prozorje in Croatia. Based on the burial characteristic, the individual is presumed to be a member of the clergy. Using stable carbon and nitrogen isotope analysis of the dentine increments from the second permanent mandibular molar (Man 2) of the individual, this study aims to identify dietary changes during the juvenile years of the individual in order to reconstruct the timing of his admission into the seminary. Mean *δ* ^13^C and *δ* ^15^N values indicate mixed C3 and C4 terrestrial based diet with the visible animal protein intake. Results of the incremental dentine analysis provide more detailed information. The possible end of the weaning period is identified, followed by the diet based on C4 resources with low animal protein intake with periods of physiological stress during his early childhood. By the end of the observed period, significant dietary change is recorded characterised by the high animal protein intake and consumption of less C4 resources. According to the historical resources, this corresponds with the timing clergy cadets usually entered the seminary with the dietary regiment consisting of daily intake of meat or fish dishes accompanied by variety of fruit and vegetables (C3 resources). This is the first such study conducted in this part of Europe and it significantly contributes to our knowledge of dietary practices and certain social customs in Early Modern Period Croatia.

## 1. Introduction

The stable isotope ratios of carbon (*δ* ^13^C) and nitrogen (*δ* ^15^N) in bone and dentine collagen have been frequently used to estimate human paleodiet. Unlike bones, which constantly remodel and represent the diet in the five to 30 years of life depending on the selected element or the age of the deceased, the primary dentine tissue does not remodel which allows us to reconstruct diet during juvenile years of the individual’s life, i.e. during the time of its formation (Nanci, 2012). Recent advancements in incremental dentine analysis enabled much higher temporal resolution (Fuller et al., 2003; Eerkens et al., 2011, Beaumont et al., 2013). Since the timing of human dentition eruption and development is well established (AlQahtani et al., 2010), microsampling of primary dentine provides a unique opportunity to track dietary changes throughout tooth’s development and recognize weaning practices (Ganiatsou et al., 2022; Velte et al., 2023), short-term dietary changes (Sánchez-Cañadillas et al. 2023), nutritional deprivation or physiological stress (Brickley et al., 2020; Nicholls et al., 2020; Feuillâtre et al., 2022).

This case study is the first time that incremental dentine analysis was performed on an archaeological sample from Croatia but also from a wider region, and aims to present a reconstruction of specific events in the early life of male individual form the church of St. Martin at Prozorje. Based on the archaeological context, the individual is presumed to have been a member of the clergy. By examining variations in *δ* ^13^C and *δ* ^15^N values of dentine sections, alongside bioarchaeological data from the skeleton and historical records, the study identifies periods of nutritional stress and dietary change, which could correspond to the individual’s admission to the seminary.

## 2. Materials

### 2.1 Archaeological site

The old ruinous parish church in the town of Dugo Selo (north-western Croatia; Fig. 1) is located at Prozorje, at the hilltop popularly called Martin’s Hill (*Martin-Breg* in Croatian), after the church’s patron saint. ‘The Land of St. Martin’ was first mentioned in a document dating back to 1209. The document was issued by Andrew II, king of Hungary and Croatia, and in it he confirms that the land in question belonged to the Knights Templar (Dobronić, 2002). Since the land was named after the saint, it implies that the church was already present at this position at the beginning of 13^th^ century, but the oldest still preserved and visible parts of the sacral building can be dated to the second half of the 15^th^ century (Belaj, 2007). The church of St. Martin at Prozorje was in continuous use from its erection until the very end of the 19^th^ century (Košćak, 2009). Except for a few minor repairs during the 20^th^ century, a large part of the abandoned building soon collapsed. In a period from 2002 to 2018, a series of archaeological excavations were conducted at the site. Complete interior of the old church was excavated along with the two old sacristies and two lateral chapels. A narrow space along some of the outside walls, primarily those along the sanctuary, was also studied during the excavation. In total, almost 300 graves were documented, the majority of which can be dated to the 17^th^ and 18^th^ centuries (Belaj and Stingl, 2019).

**Fig. 1.**
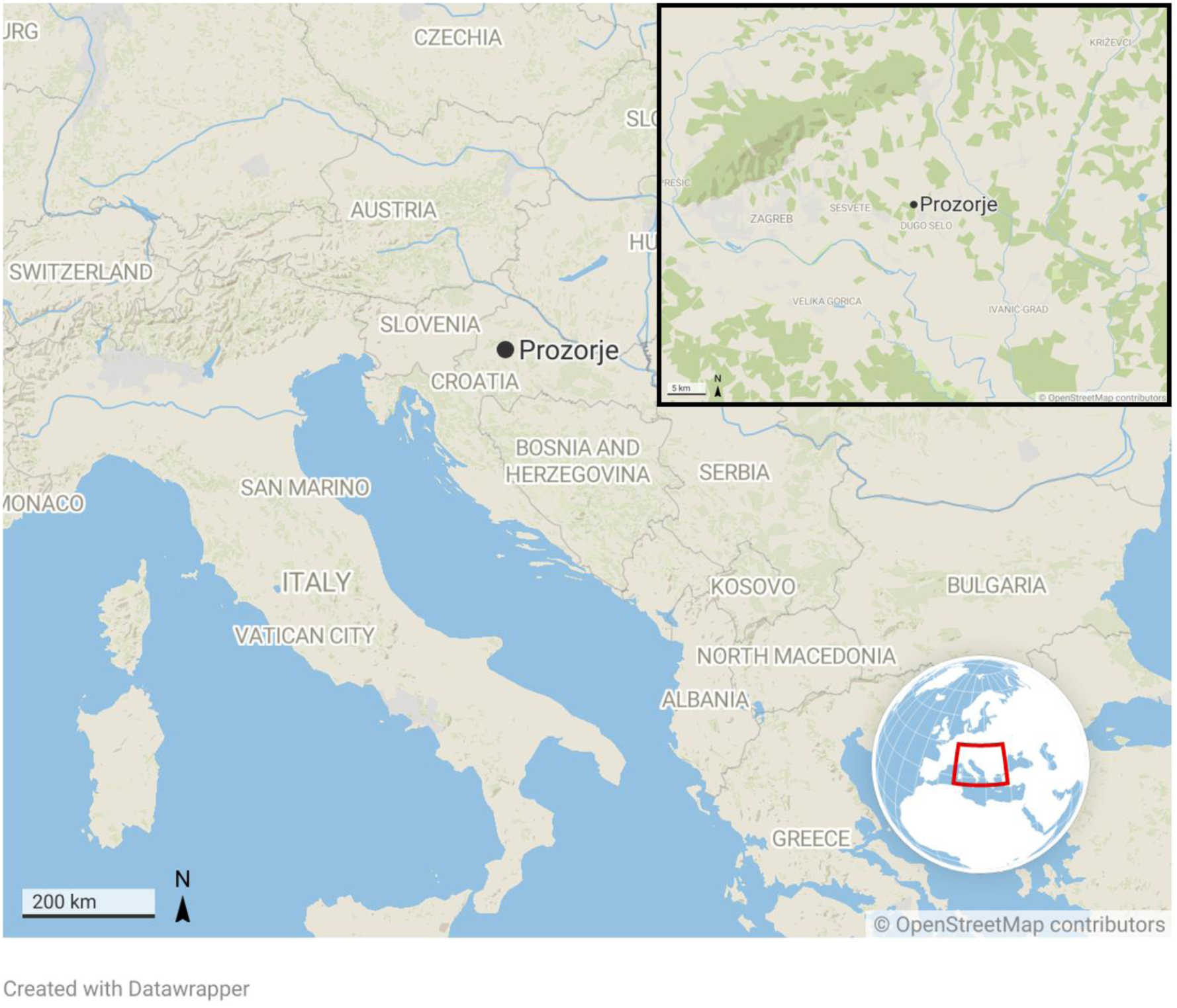
The map showing the location of the church of St. Martin at Prozorje (made by: T. Kokotović, 2024; source: OpenStreetMap, Creative Commons Attribution-ShareAlike 2.0 license, CC BY-SA 2.0; created with Datawrapper)

### 2.2 Grave no. 4

The grave no. 4 (G4) was discovered in 2002 during the first archaeological campaign. It was located in the sanctuary in front of the altar (Figs. 2 and 3) and just a little bit south of the grave 2 that was covered with a secondarily used tomb slab from the period of military orders (Belaj, 2007) which was placed on the most prominent position of the entire interior, at the central axis of the church. Poorly preserved remains of leather footwear were situated near both feet. The footwear was without iron fittings and buckles and most probably made with wooden fittings and tied with a lace. Along the individual’s forearms and on their chest, there were rows of still connected double looped hook-and-eye fasteners. Such fasteners first appeared in the territory of present-day Croatia at the end of the first half of the 14^th^ century (Anzulović, 2007) and in the archaeological context, they can be found in the layers dated from the second half of the 15^th^ century onwards (Predovnik et al., 2008). Their use rapidly spread in the 16^th^ and 17^th^ centuries (Demo, 2007). Depending on the type, they were used for hooking up shirts or for coats and socks (Azinović Bebek, 2009). Traces of wood and numerous finds of nails suggest that the deceased was buried in a coffin. Unfortunately, these finds cannot be used for a more precise dating of the grave. Given that this burial cut through all the preserved floors of the church and judging by the artefacts discovered in the two older burials found beneath, G4 can be dated to the 18^th^ century, probably to its second half (Belaj, 2006; Stingl, 2024).

**Fig. 2.**
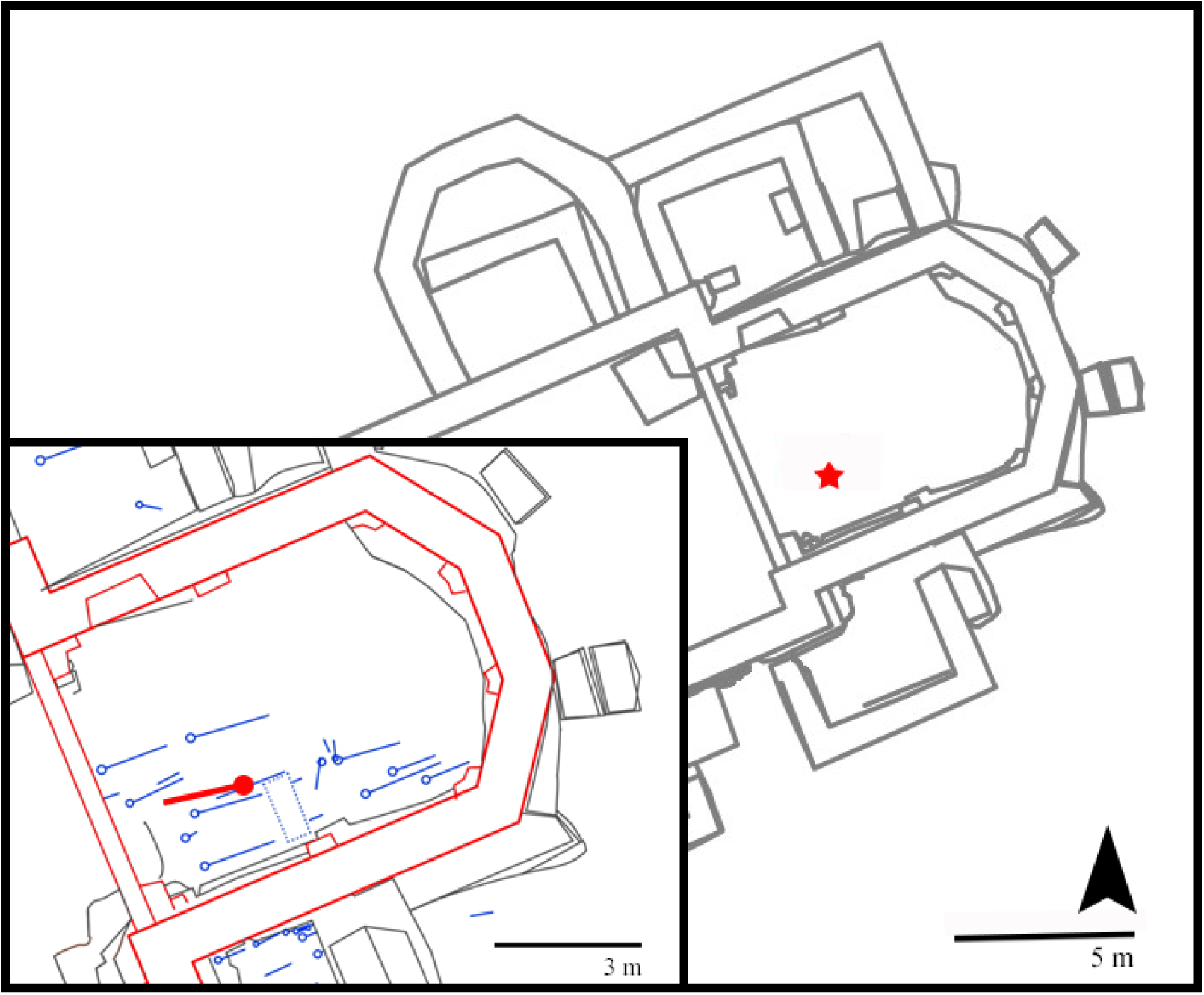
Layout of the Church of St. Martin with the marked position and orientation of G4 in the sanctuary (made by: A. Kudelić; adapted by: T. Kokotović, 2024)

**Fig. 3.**
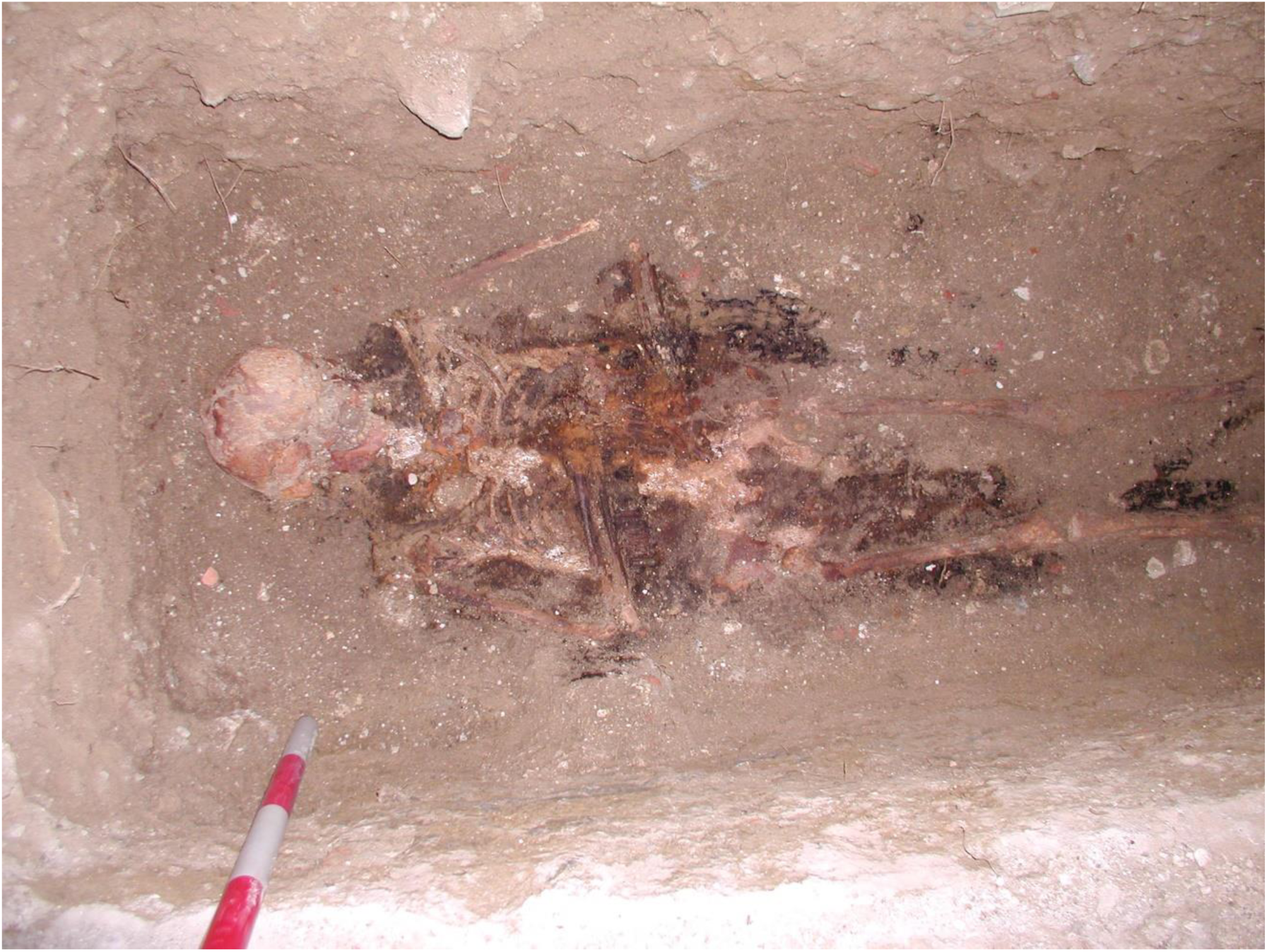
G4 during the excavation (source: Archive of the Institute of Archaeology)

What makes this burial different from the vast majority discovered at the site is its orientation. The orientation of the church of St. Martin at Prozorje is slightly deflected towards the SW-NE direction (Belaj, 2007) and most of the burials have the same orientation as the building. To simplify, since the deflection is not that significant, we can consider the orientation of the burials as predominantly west-east, which is the most common orientation of bodies in the late medieval and post-medieval periods. When the body (with or without the coffin) was laid in the grave, the head was in the west and feet in the east. However, this individual was laid in the opposite direction, with their head in the east and the feet in the west.

## 3. Methods

The sex and age-at-death were established using standard macroscopic methods and criteria (Buikstra and Ubelaker, 1994). Possible pathological changes were diagnosed based on the criteria presented in Aufderheide and Rodríguez-Martín (1998), and Ortner (2003).

For establishing approximate age of the formation of linear enamel hypoplasia (LEH), total crown height and CEJ-LEH^1^ distance was measured for each line (Buikstra, Ubleaker, 1994) using digital calliper. Approximate age of the onset of LEH on the canines was calculated using the calculator presented by Dąbrowski et al. (2021) and the method based on datasets for medieval Northern European populations (Reid and Dean 2006). This method was deemed best when considering historical and geographic context of our sample.

### 3.1. Faunal bulk C/N stable isotope analysis

The study includes faunal material from three cows (*bos taurus*) and three pigs (*sus scrofa)* from the burg Vrbovec, a medieval site adjacent to the Martin-breg. The samples are dated in the 13^th^ and 14^th^ centuries and represent the faunal baseline from carbon and nitrogen isotope ratios in the area. These are the most recent and geographically closest faunal baseline samples we were able to obtain. Collagen extraction was carried out following the modified Longin method (Longin, 1971; Brown et al., 1988). Faunal samples were demineralized in 0.5 HCl at 4°C, then placed in a HCl solution (pH3) at 70°C for 48 hours to gelatinise. Samples were then filtered using E-zee filters and freeze dried. The collagen from faunal samples was weighed out, and the samples were then sent to the Iso-Analytical Laboratory (Crewe, Cheshire, UK).

The results are reported in the conventional *δ* notation in the per mil (‰) relative to the international reported standards, VPDB for carbon, and AIR for nitrogen. The collagen quality of each sample was evaluated through the C/N atomic ratio and carbon and nitrogen content.

### 3.2. Incremental dentine analysis

Stable carbon and nitrogen isotope analysis was carried out on the incremental dentine collagen from the second permanent mandibular molar (Man 2) of the studied individual. Following Method 2 from Beaumont et al. (2013), 13 incremental dentine samples were taken from the crown to the apex of the second permanent mandibular molar (Man M2) of the individual. The tooth was first demineralized in 0.5 HCl at 4°C. Once the reaction was complete, the demineralized dentine was divided into transverse increments using a sterile scalpel, starting from the coronal dentine horn. In order for each section to produce sufficient collagen for duplicate measurements, a minimum weight after demineralization of 2.5 mg was required. The first 10 sections were divided into transverse samples at 1 mm intervals. To meet the required minimum weight of each sample, the last two increments were divided into thicker samples at 2 mm, and the final sample being divided at 5 mm. The demineralised sections were placed in a HCl solution (pH 3) at 70°C for 24 hours to gelatinise. The solution was then divided into three test tubes per sample without filtration, frozen, then freeze-dried. The collagen from human dentine samples was sent to the Isotope Ratio Mass Spectrometry Lab (SILVER) at the University of Vienna (Austria).

The approximate age of each increment was based on the timing of tooth formation described in the London Atlas (AlQahtani et al. 2010) for the permanent mandibular second molar. The method used in this study for calculating the approximate chronological age for each increment was adapted from Beaumont and Montgomery (2015). The approximate age at which mandibular second molars start to develop is around 2.5 years, and the age at root completion is 15.5 years (AlQahtani et al., 2010). The time it takes the tooth to form is approximately 13 years (15.5–2.5). This is then divided by the number of increments, in this case 19, considering that the last three increments in this case vary in thickness (10+2+2+5mm). Starting from the first coronal increment, this number (13/19∼0.7) is then added to the approximate age assigned for each increment. The approximate age for the last three increments was calculated by multiplying 0.7 with thickness of the increment (2 or 5mm).

Statistical analysis was performed in *jamovi* (version 2.5) computer software for Windows (The jamovi project (2024), retrieved from https://www.jamovi.org). The statistical test used on the data was an independent Mann-Whitney *U* test.

## 4. Results

The poorly preserved remains of a complete skeleton were discovered in the G4. Bioarchaeological analysis established the individual to be a male, aged between 30 and 45 years (middle-aged adult). The teeth of this individual show evidence of caries, alveolar abscess and LEH on the maxillary and mandibular canines. Several pathological changes (slight resorption of the bone on the inferior and lateral margins of the nasal cavity, inflammatory pitting on the palatine, and porosity around the orbits) could be indicative of leprosy (Møller-Christensen, 1961), but without additional changes to the postcranial skeleton and pathogen DNA analysis, at the moment this diagnosis is only speculative.

The total crown height and CEJ-LEH distance for both canines, as well as the estimated approximate age of onset of LEH, are presented in Tab. 1. Three lines were observed on each canine and the age of onset of LEH was estimated at around 2.5 years, then again around three and four years of age.

**Tab. 1.** Approximate age of the onset of LEH for maxillary and mandibular permanent canine of the male from grave no. 4 (made by: T. Kokotović, 2024)

|  |  | Approximate age of the onset of LEH* |
| --- | --- | --- |
| Maxillary canine |  |  |
| Total crown height/mm | 11.2 |  |
| CEJ – LEH 1 | 4.4 | 3.3 |
| CEJ – LEH 2 | 2.6 | 4.1 |
| CEJ – LEH 3 | 2.0 | 4.3 |
| Mandibular canine |  |  |
| Total crown height | 11.4 |  |
| CEJ – LEH 1 | 6.7 | 2.5 |
| CEJ – LEH 2 | 5.6 | 2.9 |
| CEJ – LEH 3 | 3.2 | 4.0 |

\*Dąbrowski et al. 2020; based on Reid, Dean 2006

The results of the faunal baseline and incremental collagen analysis are shown in Tab. 2. After the preparation of samples, all of the samples expect one (MB-4-B 10) had yielded sufficient collagen for the analysis. When human *δ*^13^C and *δ*^15^N values are compared with the faunal baseline, the expected shift in tropic level between herbivores and humans (2–5‰) is present (Ambrose 1991; Hedges and Reynard 2007). Estimated approximate age for each increment is presented in the Tab. 3.

**Tab. 2.** Isotopic data and collagen quality indicators for faunal baseline (burg Vrbovec site) and dentine sections for MB-4-B, second mandibular permanent molar (Prozorje site) (made by: T. Kokotović, 2024)

| Taxa | Element | Sample Code | Nitrogen Content % | $\delta^{15}\text{N}_{\text{AIR}}$ ‰ | Carbon Content % | $\delta^{13}\text{C}_{\text{V-PDB}}$ ‰ | C/N Ratio |
| --- | --- | --- | --- | --- | --- | --- | --- |
| bos taurus | humerus | PGV-G-A | 15.7 | 5.1 | 41.9 | −19.9 | 3.1 |
| bos taurus | radius | PGV-G-B | 15.8 | 6.5 | 42.5 | −20.3 | 3.1 |
| bos taurus | radius | PGV-G-C | 14.7 | 4.4 | 39.8 | −21.2 | 3.2 |
|  |  | **Mean** |  | 5.3 |  | −20.5 |  |
|  |  | **St. dev.** |  | 1.0 |  | 0.7 |  |
| sus scrofa | tibia | PGV-S-A | 16.1 | 6.5 | 43.4 | −20.0 | 3.2 |
| sus scrofa | radius | PGV-S-B | 16.4 | 5.1 | 44.2 | −19.8 | 3.2 |
| sus scrofa | radius | PGV-S-C | 16.1 | 6.2 | 43.6 | −21.0 | 3.2 |
|  |  | **Mean** |  | 5.9 |  | −20.3 |  |
|  |  | **St. dev.** |  | 0.7 |  | 0.7 |  |
| human | M<sub>2</sub> | MB-4-B1 | 13.6 | 9.3 | 40.0 | −14.5 | 3.4 |
|  |  | MB-4-B2 | 14.2 | 8.9 | 41.8 | −14.4 | 3.4 |
|  |  | MB-4-B3 | 14.4 | 9.4 | 42.2 | −14.1 | 3.4 |
|  |  | MB-4-B4 | 14.0 | 9.2 | 41.4 | −14.7 | 3.4 |
|  |  | MB-4-B5 | 14.4 | 8.9 | 41.8 | −15.6 | 3.4 |
|  |  | MB-4-B6 | 14.7 | 8.3 | 42.2 | −15.1 | 3.4 |
|  |  | MB-4-B7 | 14.0 | 8.8 | 41.6 | −15.2 | 3.5 |
|  |  | MB-4-B8 | 14.0 | 8.6 | 41.4 | −14.8 | 3.5 |
|  |  | MB-4-B9 | 13.2 | 9.1 | 40.0 | −14.7 | 3.5 |
|  |  | MB-4-B10 | - | - | - | - | - |
|  |  | MB-4-B11 | 13.5 | 9.6 | 39.9 | −14.6 | 3.5 |
|  |  | MB-4-B12 | 13.4 | 10.1 | 39.3 | −15.7 | 3.4 |
|  |  | MB-4-B13 | 13.7 | 11.4 | 40.6 | −17.0 | 3.5 |
|  |  | **Mean** |  | 9.3 |  | −15.0 |  |
|  |  | **St. dev.** |  | 0.8 |  | 0.8 |  |

**Tab 3.**
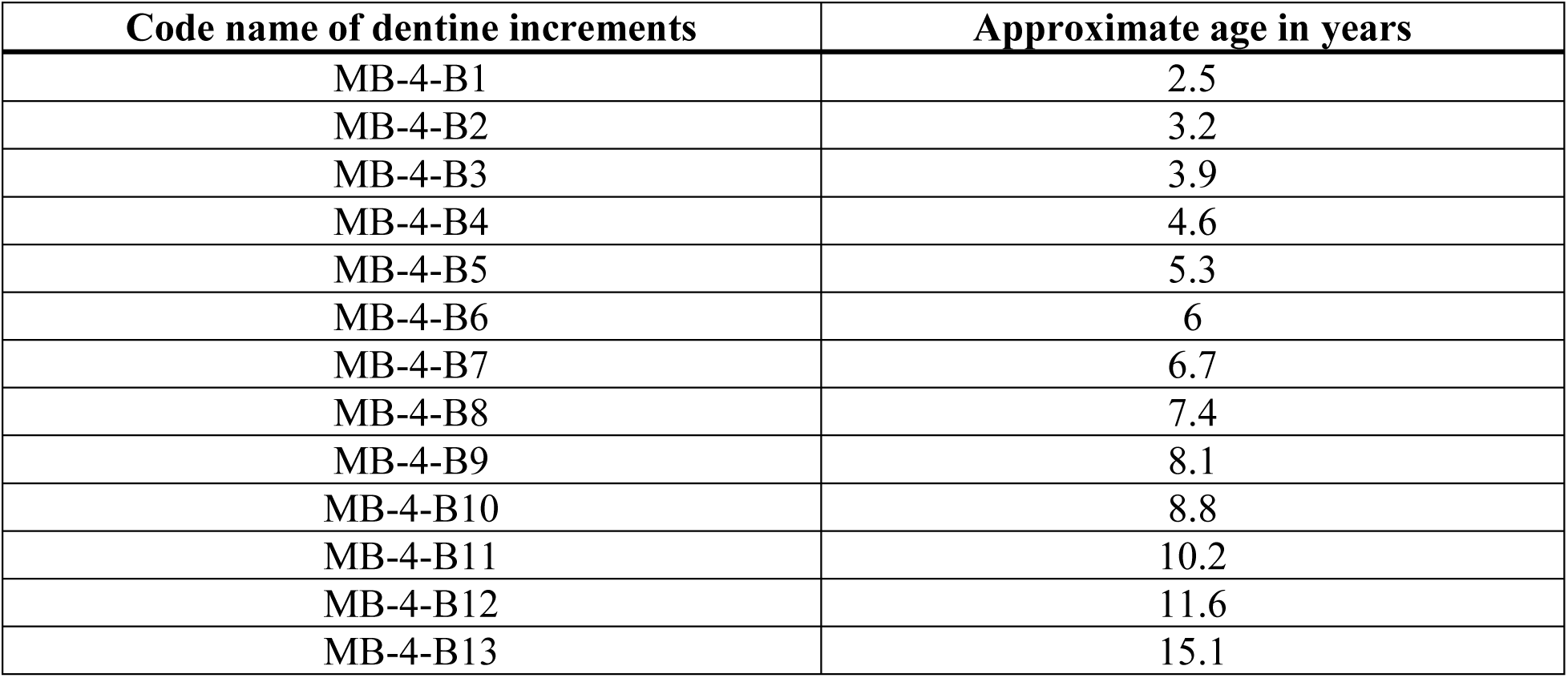
Approximate age of dentine sections for MB-4-B, second mandibular permanent molar (made by: T. Kokotović, 2024)

| **Code name of dentine increments** | **Approximate age in years** |
| --- | --- |
| MB-4-B1 | 2.5 |
| MB-4-B2 | 3.2 |
| MB-4-B3 | 3.9 |
| MB-4-B4 | 4.6 |
| MB-4-B5 | 5.3 |
| MB-4-B6 | 6 |
| MB-4-B7 | 6.7 |
| MB-4-B8 | 7.4 |
| MB-4-B9 | 8.1 |
| MB-4-B10 | 8.8 |
| MB-4-B11 | 10.2 |
| MB-4-B12 | 11.6 |
| MB-4-B13 | 15.1 |

The mean isotopic values for the second mandibular molar are –15.0‰ for *δ*^13^C and 9.3‰ for *δ*^15^N (Fig. 4). Throughout the development of the tooth, figures display a fluctuation in both *δ*^13^C and *δ*^15^N values (Figs. 5A, 5B and 6). Up to the four years of age, a slight, gradual increase is observed in *δ*^13^C (+0.4‰) values, followed by a decline (–1.5‰) around the age of six years. Between 10 and 12 years of age there is another gradual increase (+0.96‰) after which the *δ*^13^C values visibly decline (–2,4‰). *δ*^15^N values also fluctuate in the first 10 years of life, with an elevation in *δ*^15^N values (∼0.5‰) up to the age of six years, after which the values gradually increase until the age of approximately 15 years, when the recorded *δ*^15^N values are at their highest (11.45‰). The last two samples (MB-4-B-12 and MB-4-B-13), representing the period between the approximate ages of 11 and 15 years, significantly differ in both *δ*^13^C and *δ*^15^N values from the former period (independent sample Mann-Whitney *U* test p = 0.041 for both *δ*^13^C and *δ*^15^N values).

**Fig. 4.**
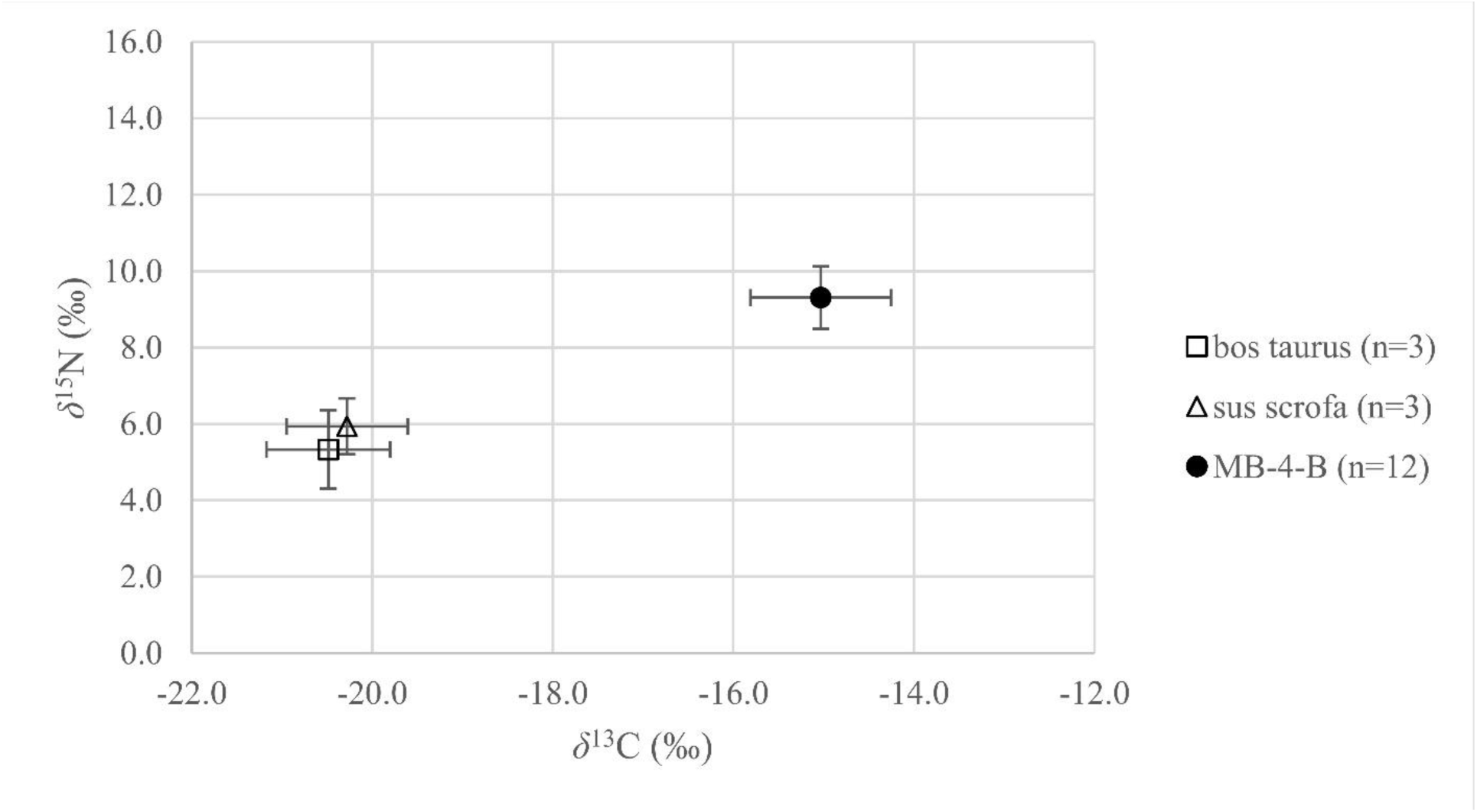
Mean *δ*^13^C and *δ*^15^N values for MB-4-B, second mandibular permanent molar (made by: T. Kokotović, 2024)

**Fig. 5A.**
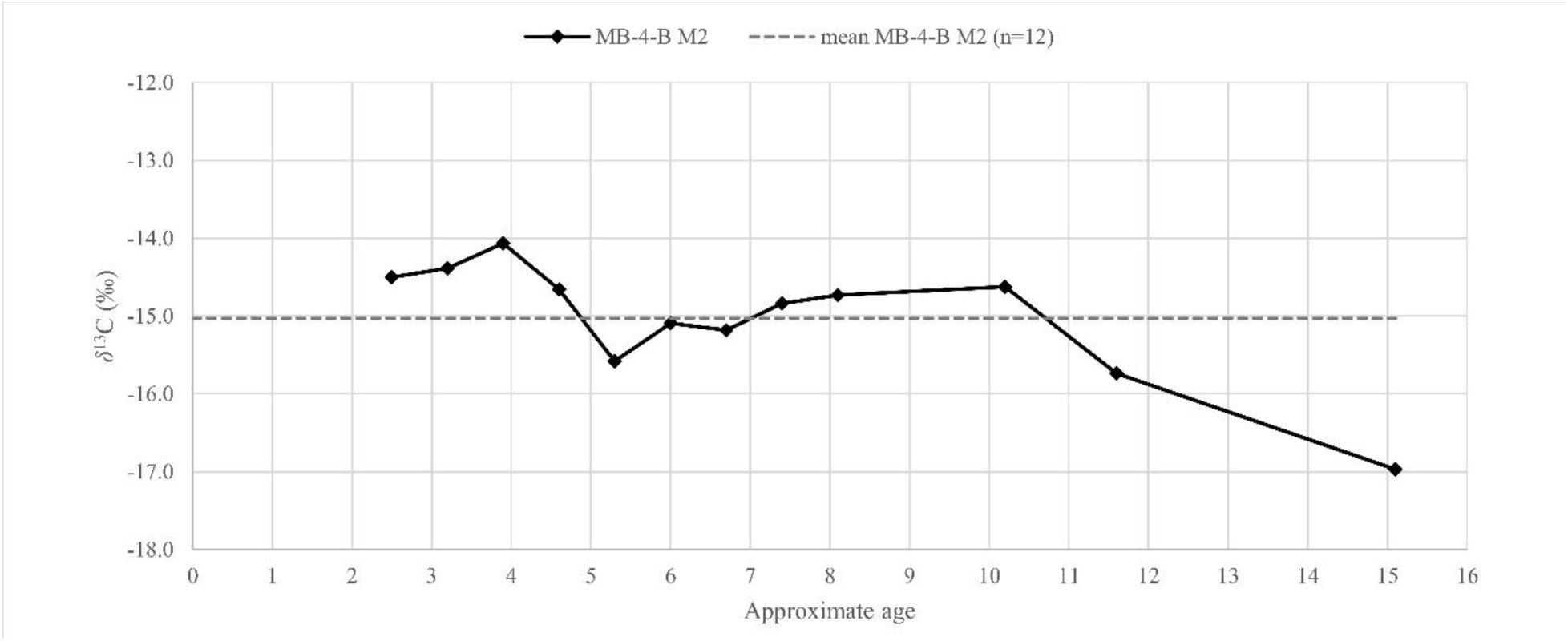
*δ*^13^C values of dentine sections against approximate age for MB-4-B, second mandibular permanent molar (made by: T. Kokotović, 2024)

**Fig. 5B.**
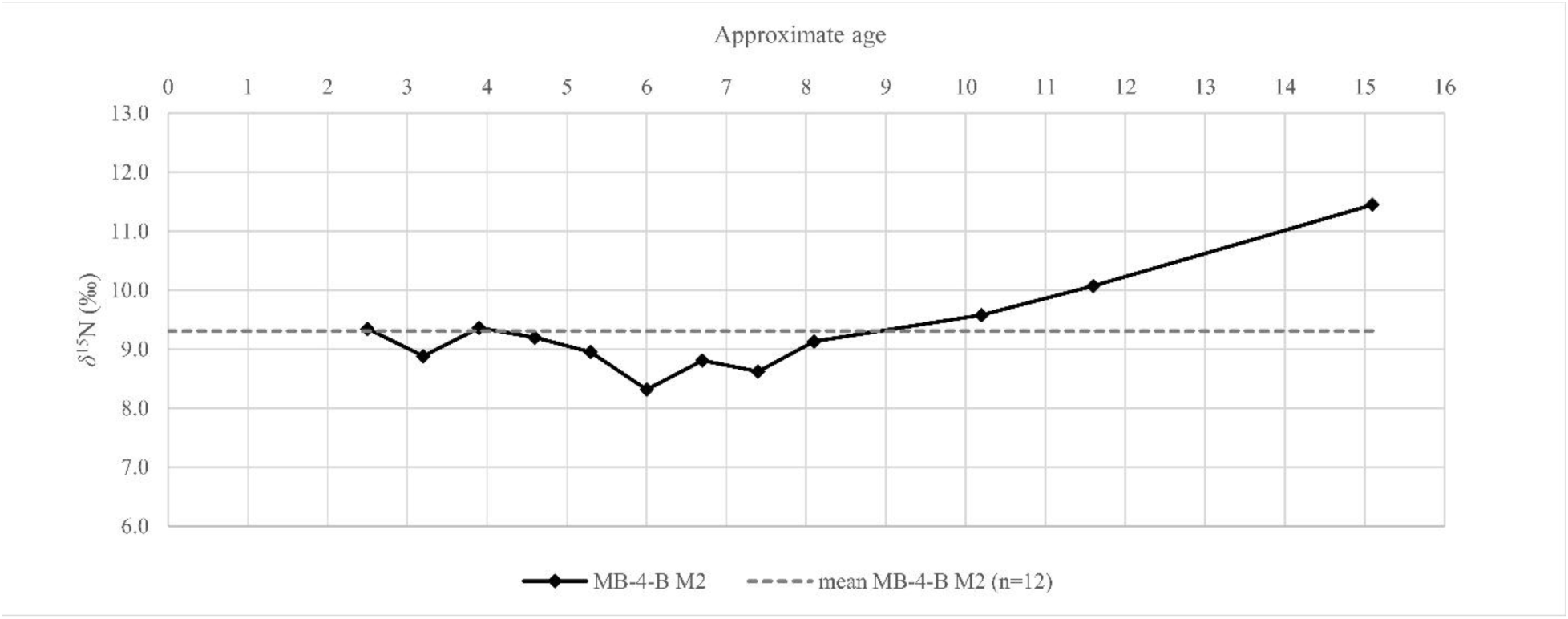
*δ*^15^N values of dentine sections against approximate age for MB-4-B, second mandibular permanent molar (made by: T. Kokotović, 2024)

**Fig. 6.**
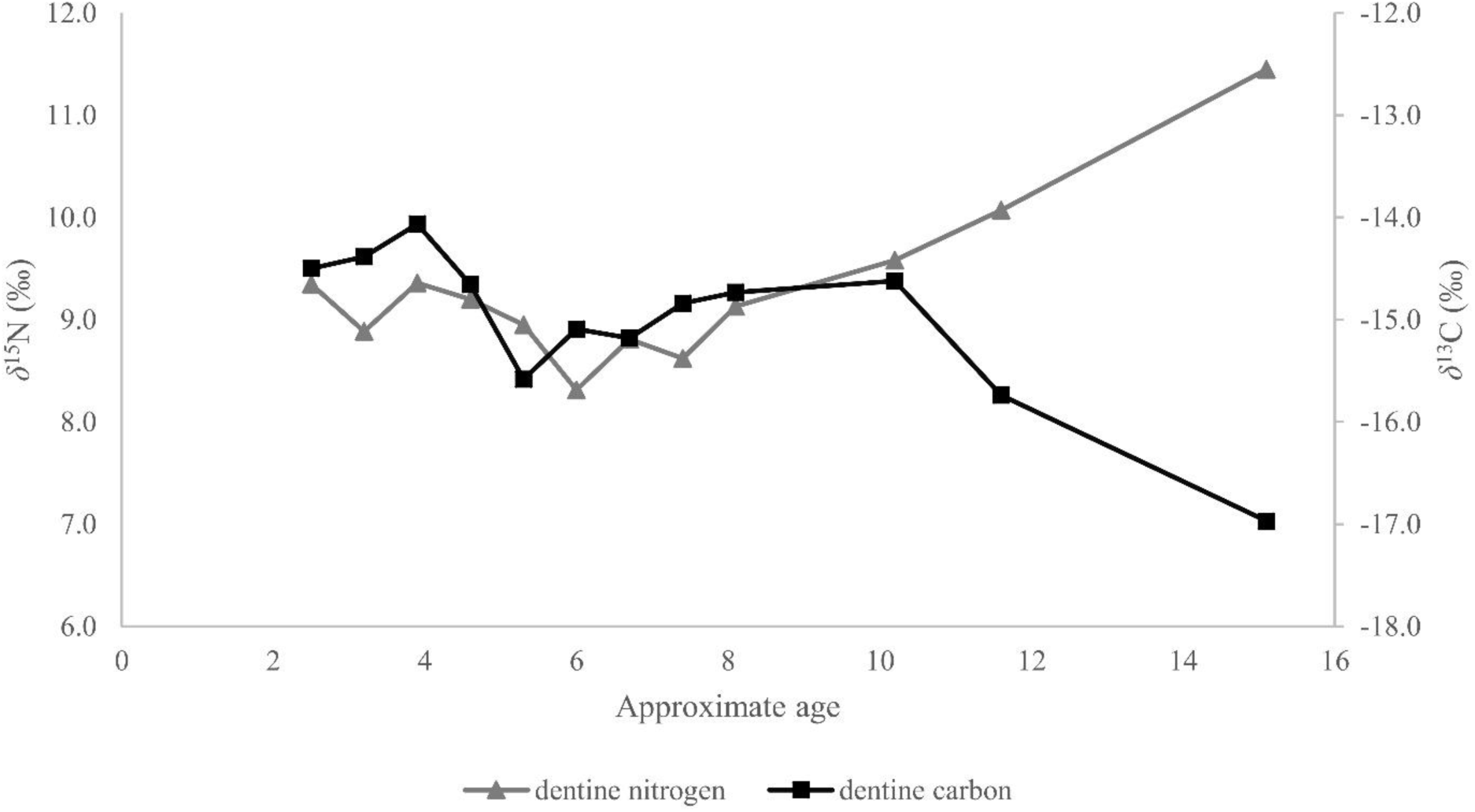
*δ*^13^C and *δ*^15^N values of dentine sections against approximate age for MB-4-B, second mandibular permanent molar (made by: T. Kokotović, 2024)

## 5. Discussion

The mean *δ*^13^C and *δ*^15^N values for the individual from G4 indicate a terrestrial staple diet based on plant food with visible dairy/animal protein intake while *δ*^13^C values indicate a mixed diet of both C3 and C4 plants. Humans consuming a diet based on C3 plants have *δ*^13^C values of ∼19‰, the ones consuming a diet consisting only of C4 plants have *δ*^13^C values of ∼ 8‰, and individuals who consume a mixed diet of both C3 and C4 plants have *δ*^13^C values that are intermediate (Dupras and Tocheri, 2007), as is the case here. These values correspond with the historical records from the 17^th^ and 18^th^ centuries about agricultural and husbandry activities in the region (Adamček, 1981). C3 plants cultivated in Dugo Selo and surrounding regions were mainly cereals (wheat, rye, oats, barley, and pyrethrum) and vines (Adamček, 1981). C4 plants included primarily millet, but also corn, whose cultivation started in the 18^th^ century (Adamček, 1981). The animals that were raised in the region were mainly pigs, chickens, cows, and horses, with primary source of animal protein being pigs, cows, and chickens, while horses and oxen were used as draft animals (Adamček, 1981).

Since the second mandibular molar starts to develop around the age of 2.5 years, exclusive breastfeeding and the start of the weaning period could not be recorded. Nevertheless, the difference in *δ* ^15^N values between the first and second samples could represent the end of the weaning process. Weaning is the period of transition from an exclusively breastmilk diet to diet combined with breastmilk, supplementary food, and drinking water, and ends with the cessation of breastfeeding (Katzenberg et al., 1996). The replacement of protein from breastmilk with protein from other foods, which are generally at a lower trophic level in respect to breastmilk, results in a drop in the infant’s *δ* ^15^N values (Fogel et al., 1989). Unfortunately, there is no specific information regarding the age of cessation of breastfeeding and supplementary foods for this region in the 17^th^ or 18^th^ centuries, but ethnographic research into the rural communities in Croatia in the 19^th^ century reveals that the duration of breastfeeding was somewhat long, from one year of age up to the birth of the second child, meaning up to two or three years on average (Čapo-Žmegač, 1998). Besides breast milk, during the weaning period, children were given easily digestible foods in the form of various porridges, made from flour, bread, or grains, especially millet (a C4 plant), diluted in milk, meat stock or water (Pleše, 2019). The nutritional content of the supplementary foods varied, and subsidizing milk and stock with water would induce various metabolic disorders (scurvy, rickets) and kidney stones, and contamination of utensils and food would result in brucellosis and/or parasitic infections (Castilho and Barros Filho, 2010). Therefore, the appearance of first and second LEH on mandibular canine (approximate age of the onset is at 2.5, 2.9 and 3.3 years) could be the result of the physiological stress connected to the weaning period. Several studies researching the connection between LEH and the weaning process observed the occurrence of LEH at the end or shortly after the end of complementary feeding (Moggi-Cecchi et al., 1994; Sandberg et al., 2014; Crowder et al., 2019; Orellana-González et al., 2020).

Another drop in the nitrogen and carbon values was recorded in the period between four and eight years, accompanied by the occurrence of LEH at the beginning of the observed period. The carbon and nitrogen values indicate terrestrial based diet with significant contribution of C4 resources and low animal/dairy protein intake, and in concurrence with LEH, could indicate a period of notable physiological stress. High levels of subadult stress were observed in several Early Modern Period populations from continental Croatia (Novak et al., 2009, 2021; Novak and Krznar, 2010; Krznar and Hajdu, 2020; Bedić et al., 2022), especially in children aged between 2 and 10 years (Novak et al., 2009; Bedić et al., 2022). Novak and colleagues (2009) suggested that the observed physiological stress was caused by a number of various factors (parasitic infections, metabolic disorders, infectious diseases) occurring as a result of inadequate nutrition, reflecting generally poor living conditions, low health and sanitary standards in continental Croatia during this period).

In our case notable changes in *δ*^13^C and *δ*^15^N values were observed after the age of 11 years: *δ*^13^C values are falling and *δ^1^*^5^N values are rising. This pattern of opposing covariance, i.e. famine pattern, was recognized by Beaumont and Montgomery (2016) when studying the victims of the Great Irish Famine and interpreted it as the evidence for the period of nutritional stress prior to their death or the introduction of famine relief food. Although this explanation cannot be completely disregarded in this case, here we see gradual change over the course of approximately five years that could also indicate visible dietary change, reflecting consumption of less C4 plants and an increase in dairy/animal protein intake during the last five and half years, but especially the last two and half years of the observed period. This shift could be related to the major lifestyle change or even change in geographical location with the access to the different food resources.

The location of the burial in the sanctuary of the church in front of the main altar, strongly suggests that the body was deliberately oriented east-west. The possibility that this orientation is a mistake caused by haste, as sometimes was the case (Krznar, 2012), can be ruled out. A most likely explanation in this case is that this grave was the burial of a priest. After the Council of Trent (1546-1563) members of the clergy were often buried with their heads in the east facing their faithful flock (Králíková, 2007). Likewise, according to the *Roman Ritual* of 1614, close attention is to be paid when burying members of the laity inside churches or chapels – they should have their feet towards the altar, while priests are supposed to have their head towards it (Bradara, 2024). This position has its roots in the belief that, upon his second arrival, Christ will come from the east and all the dead will be facing him (Tkalčec et al., 2021). For example, priest burials with this orientation were excavated in the Minster Church in Bonn (Keller, 2000) and one was documented in the filial church of St. Bartholomew in Martkt Indersdorf (Mittelstraß, 2007), both in Germany. All the graves with an east-west orientation near the main altar in the church of St. Wenceslas in Ostrava, Czechia are assumed to belong to the clergy (Králíková, 2007). Two graves with this orientation were excavated in front of the altar in the parish church of the Assumption of Mary in Hollenburg, Austria. Both are interpreted as priest burials, and for one of those two we can say this without a doubt since most parts of the vestment of the deceased could be identified: a cassock, an alb, a cingulum, a chasuble, a stole, and a maniple, revealing that he was, indeed, a priest (Leib, 2008; Grömer, 2016).

If we presume that the individual from G4 in Prozorje was indeed a member of the clergy, could a significant change in diet reflected in changes in his N/C isotopic values reflect the time when he entered the seminary? During the Council of Trent, the Decree on the establishment of the episcopal seminaries was passed. The Decree highlights a set of rules and regulations regarding the upbringing and education of clergy, prescribing the minimum age of 12 for entering the seminary (Škreblin, 1944) which corresponds with the timing of the dietary changes observed in our sample. After the Council, episcopal seminaries were established across Christian Europe in accordance with the regulations laid out by the Decree (Škalabrin, 2006). Seminary in Zagreb was established in the second half of the 16^th^ century (Patafta, 2020), and in Varaždin in 1659 (Puhmajer, 2011). Based on the available historic sources regarding the diet of the clergy cadets, it seems that their meals included substantial quantities of meat and fish that were eaten on the regular basis. According to the regulations of the Flori’s seminary in Zadar, staff and the cadets dined together and shared the same meal (Dundović, 2020: 104). They were given three meals a day (breakfast, lunch and dinner): for breakfast, they served a piece of bread with a glass of vine every day, and some fruits and cheese once a week while lunch usually consisted of soup and one meat or fish meal, depending on the day, with the addition of fruits and cheese after the main course (Dundović, 2020). Dinner included salad or soup, after which, the fruit and cheese were again consumed (Dundović, 2020). The prescribed amount of meat and/or fish amounted to 12 ounce per day/per person. Meat consumed in this seminary included beef and lamb, but not so much veal and poultry because of its scarcity in Zadar and the surrounding region (Dundović, 2020). It has to be noted that the cadets from Flori’s seminary were members of the nobility and urban population living on the Adriatic coast, so this description cannot be literally transcribed into the geohistorical context of continental Croatia. However, in the seminary in Đakovo in eastern Croatia, established at the beginning of the 19^th^ century, cadets were given two meals: lunch at 11 AM and dinner at 6:30 PM (Sršan, 1996). The cook had to prepare four different meals for lunch and three for dinner (Sršan, 1996). The meals often contained roasted meat with a side dish, and beef had to be prepared a day earlier; food had to be well cooked and clean (Sršan, 1996).

Most bulk carbon and nitrogen isotopic studies examining the diets of monastic communities in medieval and early modern Europe have primarily focused on the dietary patterns of adult members of the clergy. Studies on incremental dentine analysis examining the dietary patterns of monastic populations are limited, making this research a valuable contribution to the existing body of knowledge on the subject. The diet of these monastic communities generally resembled that of the contemporary nobility more than that of the common population. It was predominantly based on C3 plant resources and included a high intake of animal protein, as well as freshwater and marine resources (Polet and Katzenberg, 2003; Müldner and Richards, 2005; Müldner et al., 2009; Yoder, 2012; Quintelier et al., 2014; Simčenka et al., 2020; López-Costas et al., 2021; Carić and Novak, 2023).

Several incremental dentine studies have investigated the dietary changes of medieval and postmodern monastic populations in Europe. Kancle et al. (2018) investigated dietary changes within a friar population from two medieval friaries in northern England aiming to identify shifts in diet upon individuals’ entry into the religious order. Their analysis revealed a transition from a diet primarily based on terrestrial protein during early childhood to one that incorporated more marine protein by early adulthood. While the age at which these dietary changes occurred varied among individuals, the authors concluded that these changes corresponded to the period when the individuals entered the religious order. Similar study was conducted by Pliego and Beaumont (2023) at the monastery of San Milàn de la Cogolla Yuso, Spain during the 17^th^ and 18^th^ century. Their research identified distinct dietary patterns during the formative years of the monks, prior to their entry into the religious order, with some consuming lower-status foods and others higher-status foods. Following their induction, however, the monks adopted a more homogenous dietary regimen characterized by the consumption of C4 resources and a high intake of meat (Pliego and Beaumont, 2023).

## 6. Conclusion

The incremental dentine *δ* ^13^C and *δ* ^15^N profile of the male individual from the church of St. Martin at Prozorje, Croatia reveals significant insights into the dietary patterns during juvenile years of the individual in question. The isotopic values indicate a primarily terrestrial diet, including both C3 and C4 plants, alongside a noticeable intake of animal protein at the end of the profile. The shifts in isotopic values correlate with key stages during childhood, such as the end of the weaning process, as well as periods of physiological stress, potentially caused by poor nutrition, parasitic infections, or metabolic disorders, which were common in early modern period Croatia. The isotopic evidence also suggests significant dietary change after the age of 10, characterized by a reduction in C4 plant consumption and an increase in animal protein intake, which may reflect his entry into the seminary. This is supported by the burial orientation and historical context, which suggest that the studied individual was most probably a member of the clergy, and clergy cadets were known to consume diets rich in meat and fish. The findings of this study are in accord with similar studies on monastic diets in medieval and post-medieval Europe, reinforcing the idea that entering religious life often involved a transition to a diet resembling that of the higher social classes, with a greater consumption of animal protein. Overall, this study contributes to our understanding of dietary practices in Early Modern Period Croatia and the complex relationship between food, health, and social status in this period.

## Funding

This work has been supported by the Croatian Science Foundation under the project *Development and Heritage of the Military Orders in Croatia* (milOrd) (HRZZ, IP-2019-04-5513).

## Footnotes

1 CEJ-LEH – distance between cementoenamel junction (CEJ) and the LEH

## Notes

### Competing Interest Statement

The authors have declared no competing interest.

## References cited

Adamček, J., 1981. Povijest vlastelinstva Božjakovina i okolice, Kaj. XIII(IV), 105–134.

AlQahtani, S.J., Hector, M.P., Liverside, H.M., 2010. Brief communication: The London atlas of human tooth development and eruption. Am. J. Bio. Anthropol. 142(3), 481–490. 10.1002/ajpa.21258

Ambrose, S.H., 1991. Effects of diet, climate and physiology on nitrogen isotope abundances in terrestrial foodwebs. J. Archaeol. Sci. 18(3), 293–317. 10.1016/0305-4403(91)90067-Y

Anzulović, I., 2007. Ukrasno uporabni predmeti na zadarskom području u povijesnim izvorima od 13. do konca 16. Stoljeća. Radovi Zavoda za povijesne znanosti HAZU u Zadru. 49, 239–287.

Aufderheide, A.C., Rodríguez-Martín, C., 1998. The Cambridge Encyclopedia of Human Paleopathology. Cambridge University Press, Cambridge.

Azinović Bebek, A., 2009. Novovjekovni nalazi u grobovima 17. i 18. stoljeća oko crkve sv. Nikole biskupa u Žumberku. Vjesn. Arheol. Muz. u Zagrebu. 42, 463–488.

Beaumont, J., Gledhill, A., Lee-Thorp, J., Montgomery, J., 2013. Childhood diet: A closer examination of the evidence from dental tissues using stable isotope analysis of incremental human dentine. Archaeometry. 55, 277–295. 10.1111/j.1475-4754.2012.00682.x

Beaumont, J, Montgomery, J., 2015. Oral histories: a simple method of assigning chronological age to isotopic values from human dentine collagen. Ann. Hum. Biol. 42(4), 407–414. 10.3109/03014460.2015.1045027

Beaumont, J., Montgomery, J., 2016. The Great Irish Famine: Identifying Starvation in the Tissues of Victims Using Stable Isotope Analysis of Bone and Incremental Dentine Collagen. PLoS One. 11(8), e0160065. 10.1371/journal.pone.0160065

Bedić, Ž., Belaj, J., Sirovica, F., 2022. Bioarheološka studija populacije iz Gore kraj Petrinje, Hrvatska / Bioarchaeological study of the population of Gora near Petrinja, Croatia. Pril. Inst. Arheol. Zagrebu. 39(1), 173–198. 10.33254/piaz.39.1.5

Belaj, J., 2006. Interpretiranje novovjekovnih nalaza iz grobova crkve Sv. Martina na Prozorju / Interpretation of the Modern Age finds from the graves of the church of St. Martin at Prozorje. Pril. Inst. Arheol. Zagrebu. 23, 257–294.

Belaj, J., 2007. Templari i ivanovci na Zemlji Svetoga Martina. Grad Dugo Selo, Pučko otvoreno učilište, Dugo Selo.

Belaj, J., Buikstra, S., 2019. Arheološka istraživanja crkve Sv. Martina u Prozorju 2018. godine, Ann. Inst. Archaeol. XV, 173–177.

Bradara, T. 2024, Benediktinke u Puli. Arheološko istraživanje grobnica sv. Teodora / The Benedictines in Pula. The Archaeological Investigations of Tombs at the St Theodore Church Site. Monographs and catalogues – Archaeological Museum of Istria 38 (2023). Archaeological Museum of Istria, Pula.

Brickley, M.B., Kahlon, B., D’Ortenzio, L., 2020. Using teeth as tools: Investigating the mother-infant dyad and developmental origins of health and disease hypothesis using vitamin D deficiency. Am. J. Phys. Anthropol. 171(2), 342–353. 10.1002/ajpa.23947; Erratum in: Am. J. Phys. Anthropol. 2021, 175(4), 948. 10.1002/ajpa.24292

Brown, T.A., Nelson, D.E., Vogel, J.S., Southon, J.R., 1988. Improved Collagen Extraction by Modified Longin Method. Radiocarbon. 30(2), 171–177. 10.1017/S0033822200044118

Buikstra, J.E., Ubelaker, D., 1994. Standards for data collection from human skeletal remains. Arkansas Archeological Survey, Fayetteville, Arkansas.

Carić, M., Novak, M., 2023. Rekonstrukcija prehrane benediktinskih redovnica 17. i 18. stoljeća: Analiza stabilnih izotopa ugljika i dušika ljudskih ostataka iz samostanskog kompleksa Sv. Teodora u Puli / Reconstructing the diet of 17th–18th century Benedictine nuns: Carbon and nitrogen stable isotopes analysis of human remains from the St Theodore monastic complex in Pula, in: Bradara, T. (Ed.), Benediktinke u Puli. Arheološko istraživanje grobnica crkve sv. Teodora / The Benedictines in Pula. The archaeological investigation of tombs at the St Theodore Church site, Monographs and catalogues – Archaeological Museum of Istria 38 (2023). Archaeological Museum of Istria, Pula, 231–240.

Castilho, S.D., Barros Filho, A.A., 2010. The history of infant nutrition. J. Pediatr. 86(3), 179–88. 10.2223/jped.1984

Crowder, K.D., Montgomery, J., Gröcke, D.R., Filipek, K.L., 2019. Childhood “stress” and stable isotope life histories in Transylvania. Int. J. Osteoarchaeol. 29, 644– 653. 10.1002/oa.2760

Čapo-Žmegač, J., 1998. Seoska društvenost, in: Čapo Žmegač, J., Muraj, A., Vitez, Z., Grbić, J., Belaj, V. (Eds.), Etnografija: svagdan i blagdan hrvatskoga puka. Matica Hrvatska, Zagreb, 251– 295.

Dąbrowski, P., Kulus, M.J, Furmanek, M., Paulsen, F., Grzelak, J., Domagała, Z., 2021. Estimation of age at onset of linear enamel hypoplasia. New calculation tool, description and comparison of current methods. J. Anat. 239(4), 920–931. 10.1111/joa.13462

Demo, Ž., 2007.Opatovina: tragovi prošlosti izgubljene u sadašnjosti. Rezultati arheoloških iskopavanja pred crkvom svetog Franje u Zagrebu 2002. godine. Arheološki muzej, Zagreb.

Dobronić, L., 2002. Templari i ivanovci u Hrvatskoj. Dom i svijet, Zagreb.

Dundović, Z., 2020. Florijevo sjemenište u Zadru. Prilog poznavanju njegova otvorenja i djelovanja. Croatica Christiana periodica. 44(86), 87–110.

Dupras, T.L., Tocheri, M.W., 2007. Reconstructing infant weaning histories at Roman period Kellis, Egypt using stable isotope analysis of dentition. Am. J. Phys. Anthropol. 134(1), 63–74. 10.1002/ajpa.20639

Eerkens, J.W., Berget, A.G., Bartelink, E.J., 2011. Estimating weaning and early childhood diet from serial micro-samples of dentin collagen. J. Archaeol. Sci. 38(11), 3101–3111. 10.1016/j.jas.2011.07.010

Feuillâtre, C., Beaumont, J., Elamin, F., 2022. Reproductive life histories: can incremental dentine isotope analysis identify pubertal growth, pregnancy and lactation?. Ann. Hum. Biol. 49(3-4), 171–191. 10.1080/03014460.2022.2091795

Fogel, M.L., Tuross, N., Owsley, D.W., 1989. Nitrogen isotope tracers of human lactation in modern and archaeological populations. Carnegie Institution of Washington. 88, 111–117.

Fuller, B.T., Richards, M.P., Mays, S.A., 2003. Stable carbon and nitrogen isotope variations in tooth dentine serial sections from Wharram Percy. J. Archaeol. Sci. 30(12), 1673–1684. 10.1016/S0305-4403(03)00073-6

Ganiatsou, E., Vika, E., Georgiadou, A., Protopsalti, T., Papageorgopoulou, C., 2022. Breastfeeding and Weaning in Roman Thessaloniki. An Investigation of Infant Diet based on Incremental Analysis of Human Dentine. Environ. Archaeol. 1–19. 10.1080/14614103.2022.2083925

Grömer, K., 2016. Liturgical Vestments of the 16^th^ to the 18^th^ Century in Austria. Archaeological Textiles Review. 58, 10–20.

Hedges, R.E.M., Reynard, L.M., 2007. Nitrogen isotopes and the trophic level of humans in archaeology. J. Archaeol. Sci. 34(8), 1240–1251. 10.1016/j.jas.2006.10.015

Katzenberg, M.A., Herring, D.A., Saunders, S.R., 1996. Weaning and infant mortality: Evaluating the skeletal evidence. Am. J. Phys. Anthropol. 101, 177–199. 10.1002/(SICI)1096-8644(1996)23+%3C177::AID-AJPA7%3E3.0.CO;2-2

Kancle, L., Montgomery, J., Gröcke, D.R., Caffell, A., 2018. From field to fish: Tracking changes in diet on entry to two medieval friaries in northern England. J. Archaeol. Sci. Rep. 22, 264–284. 10.1016/j.jasrep.2018.07.018

Keller, C., 2000. Die barocken Klerikerbestattungen in der Münsterkirche zu Bonn. Ein Beitrag zur Verwendung von Kelchgläsern als Grabbeigabe, in: Roth, H., Joachim, H-E. (Eds.), Certamina Archaeologica. Festschrift für Heinrich Schnitzler, Bonner Beiträge zur Vor- und Frühgeschichtlichen Archäologie 1. Universität Bonn Inst. f. Vor- u. Frühgeschichtliche Archäologie Geschichte, Bonn, 229–238.

Košćak, A., 2009. Župa sv. Martina, Dugo Selo. Društvo za povjesnicu Zagrebačke nadbiskupije “Tkalčić”, Župa Sv. Martina biskupa, Zagreb, Dugo Selo.

Králíková, M., 2007. Pohřební ritus 16.–18. století na území střední Evropy (antropologicko– archeologická studie), Panorama antropologie (biologicke - socialni – kulturni). Modulove učebni texty pro studenty antropologie a „přibuznych oborů 35. Nadace Universitas, Akademicke nakladatelstvi CERM, Brno.

Krznar, S., 2012. Arheološka slika kasnosrednjovjekovnih groblja u sjevernoj Hrvatskoj. Unpublished PhD Thesis, University of Zagreb, Zagreb.

Krznar, S., Hajdu, T., 2020. Kasnosrednjovjekovna i ranonovovjekovna populacija iz Ivankova, istočna Hrvatska: rezultati (bio)arheološke analize / Late medieval/early modern population from Ivankovo, eastern Croatia: the results of the (bio)archaeological analysis. Pril. Inst. Arheol. Zagrebu. 37, 165–194. 10.33254/piaz.37.6

Leib, S., 2008. Die archäologischen Ausgrabungen in der Pfarrkirche Mariae Himmelfahrt in Hollenburg, Stadt Krems an der Donau, Niederösterreich. Fundberichte aus Österreich. 46(2007), 405–514.

Longin, R., 1971. New method of collagen extraction for radiocarbon dating. Nature. 230, 241–242. 10.1038/230241a0

López-Costas, O., Müldner, G., Lidén, K., 2021. Biological histories of an elite: Skeletons from the Royal Chapel of Lugo Cathedral (NW Spain). Int. J. Osteoarchaeol. 31(5), 941– 956. 10.1002/oa.3011

Mittelstraß, T., 2007. Die barocken Innenbestattungen in der Filialkirche St. Bartholomäus in Markt Indersdorf. Zeitschrift für Archäologie des Mittelalters. 35, 221–258.

Moggi-Cecchi, J., Pacciani, E., Pinto-Cisternas, J., 1994. Enamel hypoplasia and age at weaning in 19th-century Florence, Italy. Am. J. Phys. Anthropol. 93(3), 299–306. 10.1002/ajpa.1330930303

Møller-Christensen, V., 1961. Bone Changes in Leprosy. Munskgaard, Copenhagen.

Müldner, G., Montgomery, J., Cook, G., Ellam, R., Gledhill, A., Lowe, C., 2009. Isotopes and individuals: diet and mobility among the medieval Bishops of Whithorn. Antiquity. 83(322), 1119–1133. 10.1017/S0003598X00099403

Müldner, G., Richards, M.P., 2005. Fast or feast: reconstructing diet in later medieval England by stable isotope analysis. J. Archaeol. Sci. 32(1), 39–48. 10.1016/j.jas.2004.05.007

Nanci, A., 2012. Ten Cate’s Oral Histology. Mosby, St. Louis.

Nicholls, R., Buckberry, J., Beaumont, J., Črešnar, M., Mason, P., Armit, I., Koon, H., 2020. A carbon and nitrogen isotopic investigation of a case of probable infantile scurvy (6th–4th centuries BC, Slovenia). J. Archaeol. Sci. Rep. 30, 102206. 10.1016/j.jasrep.2020.102206

Novak, M., Šlaus, M., Pasarić, M., 2009. Subadultni stres u srednjovjekovnim i novovjekovnim populacijama kontinentalne Hrvatske / Subadult Stress in the Medieval and Early Modern Populations of Continental Croatia. Pril. Inst. Arheol. Zagrebu. 26, 247–270.

Novak, M., Krznar, S., 2010. Antropološka analiza ljudskoga kostura s nalazišta Torčec-Prečno pole I, in: Sekelj Ivančan, T., Podravina u ranom srednjem vijeku. Rezultati arheoloških istraživanja ranosrednjovjekovnih nalazišta u Torčecu, Monographiae Instituti archaeologici 2. Institut za arheologiju, Zagreb.

Novak, M., Bedić, Ž., Los, Dž., Premužić, Z., 2021. Bioarchaeological analysis of the 15^th^ –17^th^ century population from Zrin, continental Croatia. A. Rad. Raspr. 20 (1), 317–336. 10.21857/y7v64t0wxy

Orellana-González, E, Sparacello, V.S., Bocaege, E., Varalli, A., Moggi-Cecchi, J., Dori, I., 2020. Insights on patterns of developmental disturbances from the analysis of linear enamel hypoplasia in a Neolithic sample from Liguria (northwestern Italy). Int. J. Paleopathol. 28, 123–136. 10.1016/j.ijpp.2019.12.005

Ortner, D.J., 2003. Identification of Pathological Conditions in Human Skeletal Remains. Academic Press, USA. 10.1016/B978-0-12-528628-2.X5037-6

Patafta, D., 2020. Razvoj teoloških i filozofskih studija u Zagrebu od XIII. stoljeća do osnivanja Sveučilišta u Zagrebu. Povijesni korijeni Katoličkoga bogoslovnog fakulteta. Bogosl. Smotra. 90(1), 43–66.

Pleše, I., 2019. Svaka majka doji svoje dite, samo ako ima mlika: kako čitamo Zbornik za narodni život i običaje južnih Slavena. Nar. umjet. 56(2), 57–73. 10.15176/vol56no203

Pliego, A.H., Beaumont, J., 2023. The monks of San Millán: Investigating the transition between pre-monastic and monastic diet using carbon and nitrogen isotope ratios in incremental dentine. J. Archaeol. Sci. Rep. 49, 103981. 10.1016/j.jasrep.2023.103981

Polet, C., Katzenberg, M.A., 2003. Reconstruction of the diet in a mediaeval monastic community from the coast of Belgium. J. Archaeol. Sci. 30(5), 525–533. 10.1016/S0305-4403(02)00183-8

Predovnik, K., Dacar, M., Lavrinc, M., 2008. Cerkev Sv. Jerneja v Šentjerneju: arheološka iskopavanja v letih 1985 in 1986, Archaeologia historica Slovenica 6. Univerza v Ljubljani, Filozofska fakulteta, Znanstvena založba, Oddelek za arheologijo, Ljubljana.

Puhmajer, P., 2011. Zgrada Zakmardijeva sjemeništa u Varaždinu — projekt, izgradnja, tipologija. Peristil. 54(1), 151–160.

Quintelier, K., Ervynck, A., Müldner, G., Van Neer, W., Richards, M.P., Fuller, B.T., 2014. Isotopic examination of links between diet, social differentiation, and DISH at the post-medieval Carmelite Friary of Aalst, Belgium. Am. J. Phys. Anthropol. 153(2), 203–213. 10.1002/ajpa.22420

Reid, D.J., Dean, M.C., 2006. Variation in modern human enamel formation times. J. Hum. Evol. 50(3), 329–46. 10.1016/j.jhevol.2005.09.003

Sánchez-Cañadillas, E., Beaumont, J., Santana-Cabrera, J., Gorton, M., Arnay-de-la-Rosa, M., 2023. The early lives of the islanders: Stable isotope analysis of incremental dentine collagen from the prehispanic period of the Canary Islands. Am. J. Biol. Anthropol. 182(2), 300–317. 10.1002/ajpa.24828

Sandberg, P.A., Sponheimer, M., Lee-Thorp, J., Van Gerven, D., 2014. Intra-tooth stable isotope analysis of dentine: a step toward addressing selective mortality in the reconstruction of life history in the archaeological record. Am. J. Phys. Anthropol. 155(2), 281–93. 10.1002/ajpa.22600

Simčenka, E., Jakulis, M., Kozakaitė, J, Piličiauskienė, G., Lidén, K., 2020. Isotopic dietary patterns of monks: results from stable isotope analyses of a seventeenth–eighteenth century Basilian monastic community in Vilnius, Lithuania. Archaeol. Anthropol. Sci. 12(102), 102–116. 10.1007/s12520-020-01063-9

Sršan, S., 1996. Statut Bogoslovnog sjemeništa u Đakovu. Diacovensia. 4(1), 147–166.

Stingl, S., 2024. The first among equals? – Review of the burial chambers and graves with some architectural elements inside of the Church of St Martin at Prozorje, in: Belaj, J., Hunyadi, Z., Tkalčec, T., Krznar, S., Sekelj Ivančan, T., Karavidović, T., Kokotović, T., Stingl, S. (Eds.), Military Orders and Their Heritage, Proceedings of the 8^th^ International Conference on Mediaeval Archaeology of the Institute of Archaeology, 9^th^–10^th^ November 2022, Zagreb. Institut za arheologiju, Zagreb.

Škalabrin, N., 2006. Biskupijsko sjemenište u Đakovu – od osnutka do 1918. Diacovensia. 14 (2), 215–257.

Škreblin, I., 1944. Odgoj i nastava u Zagrebačkom sjemeništu 1578. – 1900. in: Kniewald, D. (Ed.), Kulturno – poviestni Zbornik Zagrebačke nadbiskupije u spomen 850. godišnice osnutka I. Hrvatsko izdavalački bibliografski zavod, Zagreb.

Tkalčec, T., Sekelj Ivančan, T., Krznar, S., 2021. Arheologija srednjovjekovnih utvrda, naselja i groblja sjeverne Hrvatske. Monografije Instituta za arheologiju 5. Institut za arheologiju, Zagreb.

Velte, M., Czermak, A., Grigat, A., Neidich, D., Trautmann, B., Lösch, S., Päffgen, B., Harbeck, M., 2023. Tracing early life histories from Roman times to the Medieval era: weaning practices and physiological stress. Archaeol. Anthropol. Sci. 15, 190. 10.1007/s12520-023-01882-6

Yoder, C., 2012. Let them eat cake? Status-based differences in diet in medieval Denmark. J. Archaeol. Sci. 39(4), 1183–1193. 10.1016/j.jas.2011.12.029

